# Activity-regulated growth of motoneurons at the neuromuscular junction is mediated by NADPH oxidases

**DOI:** 10.1101/2022.10.27.514147

**Authors:** Daniel Sobrido-Cameán, Matthew C. W. Oswald, David M. D. Bailey, Amrita Mukherjee, Matthias Landgraf

## Abstract

Neurons respond to changes in the levels of activity they experience in a variety of ways, including structural changes at pre- and postsynaptic terminals. An essential plasticity signal required for such activity-regulated structural adjustments are reactive oxygen species (ROS). To identify sources of activity-regulated ROS required for structural plasticity *in vivo* we used the *Drosophila* larval neuromuscular junction as a highly tractable experimental model system. For adjustments of presynaptic motor terminals, we found a requirement for both NADPH oxidases, Nox and Dual Oxidase (Duox), that are encoded in the *Drosophila* genome. This contrasts with the postsynaptic dendrites from which Nox is excluded. NADPH oxidases generate ROS to the extracellular space. Here, we show that two aquaporins, Bib and Drip, are necessary ROS conduits in the presynaptic motoneuron for activity regulated, NADPH oxidase dependent changes in presynaptic motoneuron terminal growth. Our data further suggest that different aspects of neuronal activity-regulated structural changes might be regulated by different ROS sources: changes in bouton number require both NADPH oxidases, while activity-regulated changes in the number of active zones might be modulated by other sources of ROS. Overall, our results show NADPH oxidases as important enzymes for mediating activity-regulated plasticity adjustments in neurons.

## 1 Introduction

Reactive oxygen species (ROS) have commonly been associated with detrimental processes such as oxidative stress, toxicity, ageing, neurodegeneration and cell death because increases in ROS levels seen with ageing and neurodegenerative disorders, including Parkinson’s (Spina and Cohen; 1989) and Alzheimer’s disease (Martins et al; 1986). However, it is appreciated that ROS are not simply cytotoxic agents, but more generally function as signalling molecules in a multitude of processes, including growth factor signalling (Suzukawa et al., 2000; Goldsmit et al., 2001; Kamata et al., 205; Nitte et al., 2010), wound healing (Razzell et al; 2013) and in development (Milton et al., 2011; Oswald et al; 2018a; Dhawan et al., 2020; for a reviews see Owusu-Ansah and Banerjee, 2009; Massaad and Klann, 2011; Wilson and Gonzalez-Billaut, 2015; Oswald et al., 2018b; Terzi and Suter, 2020).

During nervous system development, ROS signalling is involved at all stages, from neurogenesis to pathfinding to synaptic transmission (Knapp and Klann, 2002; Kishida and Klann, 2007; Massaad and Klann, 2011; Wilson and Gonzalez-Billaut, 2015; Wilson et al., 2018; Terzi and Suter, 2020). When studying ROS signalling *in vivo*, challenges include the ability to disentangle cell autonomous from indirect or systemic effects; or to determine sources and types of ROS. Using the fruit fly, *Drosophila melanogaster*, as a highly tractable experimental model system, genetic manipulations targeted to single motoneurons were able to identify hydrogen peroxide as a synaptic plasticity signal, generated as a consequence of neuronal overactivation and both necessary and sufficient for activity-regulated adaptive changes of synaptic terminal structure and transmission (Oswald et al., 2018; Dhawan et al., 2020). We found mitochondria to be a major source of activity-regulated hydrogen peroxide with opposing effects on the growth of pre- *vs* postsynaptic terminals: at the presynaptic terminal of the neuromuscular junction (NMJ) overactivation and hydrogen peroxide cause increases in terminals (Milton et al., 2011; Oswald et al., 2018). This change in presynaptic terminal growth is mediated by activation of the JNK signalling pathway (Milton et al., 2011), and it utilises the conserved Parkinson’s disease-linked protein, DJ-1b, as a redox sensor, which regulates the PTEN-PI3 Kinase growth pathway (Oswald et al., 2018). In contrast, the size of postsynaptic dendritic arbors is negatively regulated by over-activation and activity-regulated hydrogen peroxide (Tripodi et al., 2008; Oswald et al., 2018; Dhawan et al., 2020). These studies using the *Drosophila* larval neuromuscular model system contrast with findings from cultured hippocampal neurons, which posit mitochondrially generated superoxide as the principal ROS signal downstream of over-activation (Hongpaisan et al. 2003; 2004). The extent to which both types of ROS operate as neuronal plasticity signals downstream of over-activation remains to be resolved, though it is possible that apparent discrepancies might be due to the use of different cellular models and/or a reflection of the degree of overactivation.

Another principal source of ROS are NADPH oxidases, whose location in the plasma membrane could facilitate sub-cellular signalling discrete from mitochondrial ROS production. NADPH oxidases are integral membrane proteins that mediate a single electron transfer from NADPH to oxygen, thereby converting it to superoxide (Lambeth, 2002). These enzymes are prevalent throughout the evolutionary ladder from Amoebozoa and fungi to higher plants and mammals. NADPH oxidases are involved in growth and plasticity during nervous system development (Kishida et al., 2006; Munnamalai and Suter, 2009; Munnamalai et al., 2014; Olguín-Albuerne and Morán, 2015; Serrano et al., 2003; Tejada-Simon et al., 2005; Wilson et al., 2015; Wilson et al., 2016; Terzi and Suter, 2020). In contrast to mammalian genomes, which encode seven Nox isoforms (Nox 1-5, Duox 1 and 2) (Lambeth et al., 2002; Kawahara et al., 2007), *Drosophila melanogaster* encodes just two NADPH oxidases: dual oxidase (*Duox*) and a Nox-5 homolog (*Nox*). Enzymatic activity of both is calcium-regulated, via their N-terminal calcium binding EF-hands (Razzell et al, 2013; Ha et al., 2005b, 2009; S. Moreira et al., 2010). Curiously, the mouse genome does not encode a calcium-regulated Nox-5 homologue, which has therefore not been studied extensively *in vivo* (Kawahara et al, 2004). Recently, we identified the NADPH oxidase Duox as necessary in motoneurons to reduce their dendritic arbors in response to neuronal over-activation, an adaptive response to reduce the numbers of presynaptic inputs and thus synaptic drive (Zwart et al., 2013; Dhawan et al., 2020). We further found that these activity-regulated ROS generated by Duox at the extracellular face of the plasma membrane, required the aquaporins, Bib and Drip; presumably for efficient entry into the cytoplasm to regulate dendritic growth and/or stability (Dhawan et al., 2020).

Here, we investigated the role of NADPH oxidases at the presynaptic terminal of the NMJ, whose growth response to neuronal over-activation is distinct to that of the dendritic compartment of the motoneuron. We show that the NADPH oxidases Duox and Nox are sources of activity-regulated ROS that mediate activity-regulated growth of NMJ terminals. In contrast to motoneuron dendrites, both NADPH oxidases function at the presynaptic NMJ, necessary and sufficient to elicit changes in growth. At the NMJ too, we find the aquaporins, Bib and Drip, are necessary for autocrine signalling at the NMJ. This arrangement at the presynaptic NMJ terminal contrasts with their dendritic function within these motoneurons, where only Duox, but not Nox, is required. This differential requirement of Nox mirrors its sub-cellular localisation, with Nox largely excluded from dendrites. Furthermore, at the postsynaptic compartment extracellular ROS, including from other neurons in the vicinity, act as local plasticity signals that cause reductions in dendritic arbor size (Dhawan et al., 2020).

## 2 Results

### NADPH oxidases, Duox and Nox, are both required for activity-regulated growth at the neuromuscular junction

Mitochondria are a major source of activity-generated ROS, notably within the cytoplasm. Here, we sought to investigate the role of membrane localised ROS generators, the NADPH oxidases Nox and Duox, during activity-regulated adjustment of presynaptic terminals. As a highly tractable experimental model we used the well characterised neuromuscular junction (NMJ) of the Drosophila larva (Frank et al., 2013). Specifically, we focused on the NMJ of the so called ‘anterior Corner Cell’ (aCC), which innervates the most dorsal body wall muscle, known as muscle 1 (Crossley 1978) or dorsal acute muscle 1 (DA1) (Sink and Whitington, 1991; Landgraf et al., 1997; Baines et al., 1999; Baines et al., 2001; Bate, 1993; Choi et al., 2004; Hoang and Chiba, 2001). For cell-specific over-activation of aCC motoneurons, we used the established paradigm of targeted mis-expression of the warmth-gated cation channel, dTRPA1 (Hamada et al., 2008; Oswald et al., 2018; Dhawan et al., 2020). This allows aCC motoneurons to be selectively over-activated simply by placing larvae at >24°C, the temperature threshold for dTRPA1 ion channel opening (Pulver et al., 2009).

First, we re-confirmed that at 25°C *dTrpA1* expression in aCC motoneurons leads to significant increases in bouton number at the aCC-DA1 NMJ relative to non-manipulated controls, as previously shown (Oswald et al., 2018) (Figure 1). An advantage of using cell-specific dTRPA1-mediated activity manipulations in this system is that these can be carried out at 25°C, a temperature considered optimal for *Drosophila melanogaster* development (Lachaise et al., 1988; Pool et al., 2012) and therefore generally considered neutral, while sufficient to mildly activate neurons that mis-express dTRPA1 (Pulver et al., 2009; Tsai et al., 2012).

**Figure 1.**
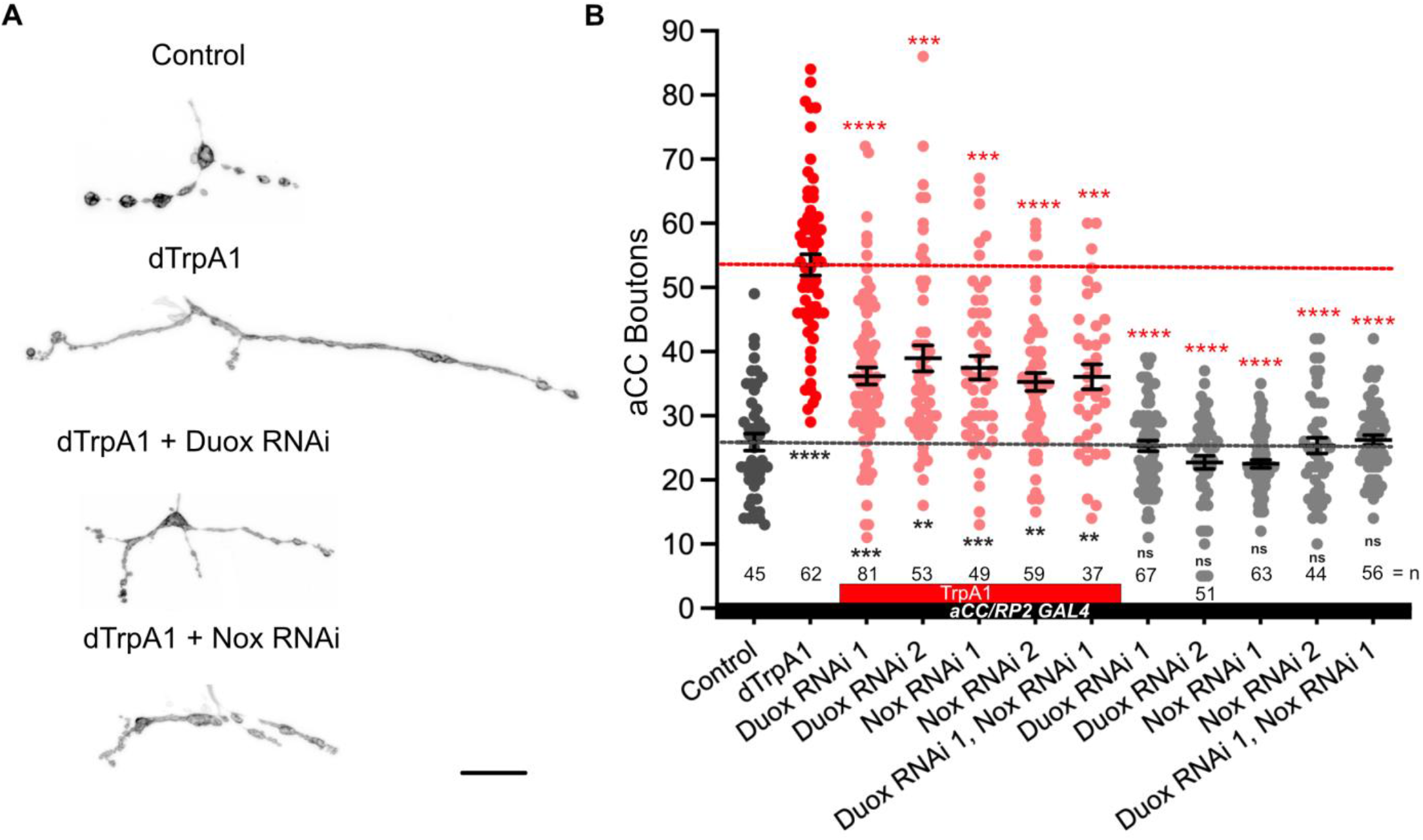
NADPH oxidases, dDuox and dNox, are both required for activity-regulated growth of the neuromuscular junction. A) Representative images of aCC motoneuron terminals on their target muscle, DA1 [muscle 1, according to (Crossley, 1978)] in third instar larvae (100 hr ALH): control; dTrpA1 overactivated; dTrpA1 overactivated while either Duox or Nox is concomitantly knocked down via targeted RNAi (“TrpA1 + Duox KD” and “TrpA1 + Nox KD”). B) Dot-plot quantification shows NMJ bouton number increases in response to cell-specific activity increases. This phenotype is rescued by simultaneous NADPH oxidase knockdown. Triangles represent presence of UAS-dTrpA1 activity manipulation, while circle indicate absence of dTrpA1. Mean ± SEM, ANOVA, ** p<0.01, *** p<0.001, **** p<0.0001. Red asterisks indicate comparisons with the UAS-TrpA1 over-activation group, while black indicate comparison with the un-manipulated wild type control. Scale bar = 20 μm.

Next, we tested the requirement for the two NADPH oxidases encoded in the Drosophila genome, Duox and Nox, in mediating these activity-regulated structural changes at the NMJ. To this end, we expressed RNAi transgenes for knocking down endogenous Duox or Nox in aCC motoneurons. By themselves, expression of *UAS-Duox-RNAi* or *UAS-Nox-RNAi* transgenes in aCC motoneurons have no measurable effect on NMJ morphology. However, in motoneurons that have been overactivated by *UAS-dTrpA1*, the characteristic activity-induced bouton overgrowth phenotype is suppressed by co-expression of *UAS-Duox-RNAi* or *UAS-Nox-RNAi* transgenes, individually or combined (Figure 1). Neuronal overactivation by *UAS-dTrpA1* also causes a reduction in active zone numbers (Oswald et al., 2018). We find no statistically significant changes in active zone number following NADPH oxidases manipulations (Supplementary figure 1). These results show that the membrane localised ROS generators, Nox and Duox, are required primarily for activity-regulated changes in presynaptic terminal growth while not significantly impacting on the number of presynaptic release sites.

### Duox and Nox activity is sufficient for mediating structural changes at the NMJ

We next asked if the activity of these NADPH oxidases might also be sufficient for regulating presynaptic terminal growth. To test this, we overexpressed *UAS-Duox* or *UAS-Nox* transgenes in aCC motoneurons. Quantification showed comparable increases in bouton number at the NMJ as a consequence of over-expression of either Duox or Nox. No enhancement of this phenotype occurs when both are co-expressed (Figure 2). In contrast, active zone numbers are not significantly impacted by overexpression of either NADPH oxidase (Supplementary figure 1).

**Figure 2.**
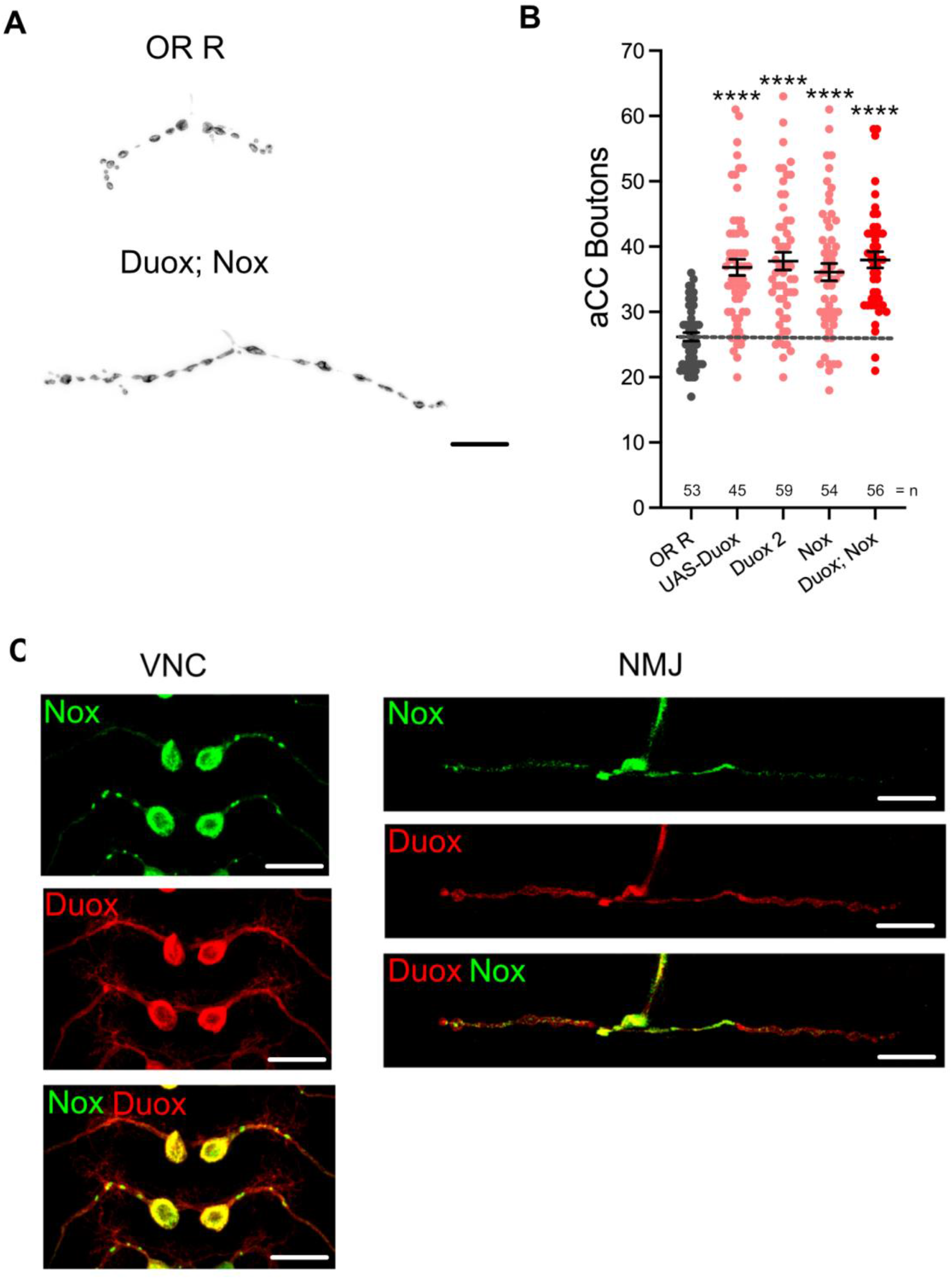
dDuox or dNox activity is sufficient for mediating structural changes at the NMJ. A) Representative images of aCC presynaptic terminals on muscle DA1 from third instar larvae (100 hr ALH) of control aCC and those overexpressing Duox and Nox. B) Dot-plot quantification shows NMJ bouton number increases in response to cell-specific over-expression of NADPH oxidases. C) Duox and Nox, localization in neurons: representative confocal micrograph images of aCC somata and dendrites in the ventral nerve cord (VNC) and aCC presynaptic terminals at the DA1 muscle in third instar larvae (72 hr ALH), showing subcellular localisation of tagged over-expressed Duox∷mRuby2∷HA (in red) and Nox∷YPet (in green). Mean ± SEM, ANOVA, **** p<0.0001. Comparisons are made with the control group. Scale bar = 20 μm.

For the postsynaptic compartment, namely the dendritic arbor of motoneurons, we had previously shown that only Duox, but not Nox, has a role in activity-regulated plasticity (Dhawan et al., 2020). To further explore this difference in NADPH oxidase requirement between pre-*vs* postsynaptic compartments, we generated tagged transgenes of both NADPH oxidases, *UAS-Duox∷mRuby2∷HA* and *UAS-Nox∷YPet*. When expressed in aCC motoneurons to reveal sub-cellular localisation, we see exclusion of Nox∷YPet from the postsynaptic dendrites, while Duox∷mRuby2∷HA is fairly homogeneously distributed within the plasma membrane (Figure 2C). These patterns of distinct sub-cellular distributions, notably exclusion of Nox∷YPet from dendrites, are compatible with the genetic manipulations phenotypes and point to Nox being selectively sorted to soma and presynaptic compartments in these neurons.

### Aquaporin channel proteins Bib and Drip are necessary for NADPH oxidase-regulated structural changes at the NMJ

The NADPH oxidases Duox and Nox are transmembrane proteins that generate ROS at the extracellular face of the plasma membrane (Lambeth, 2002; Panday et al., 2015). We reasoned that if NADPH oxidase-generated ROS are indeed instrumental in activity-regulated adjustment of synaptic terminals, then neutralisation of extracellular ROS should rescue NMJ phenotypes associated with NADPH oxidase overexpression. To test this, we mis-expressed in aCC motoneurons two different forms of catalases that are secreted to the extracellular space; a human version and the Drosophila immune-regulated catalase (Irc) (Ha et al., 2005b; Fogarty et al., 2016). These catalases neutralise extracellular hydrogen peroxide by conversion to water. On their own, their mis-expression in aCC motoneurons has no significant impact on NMJ structure or size. To test the model of neuronal activity leading to NADPH oxidase activation, leading to extracellular ROS production, we co-expressed secreted catalase in aCC motoneurons while over-activating these with dTRPA1. The presence of a secreted catalase suppresses the NMJ growth that would otherwise ensue with neuronal overactivation (Figure 3A). Similarly, NMJ over-growth stimulated by over-expression of Duox is also neutralised by co-expression of secreted catalase in the same neuron (Figure 3B). These experiments demonstrate that it is the presence of extracellular ROS, notably hydrogen peroxide generated by NADPH oxidases, which leads to activity-induced changes in NMJ growth.

**Figure 3.**
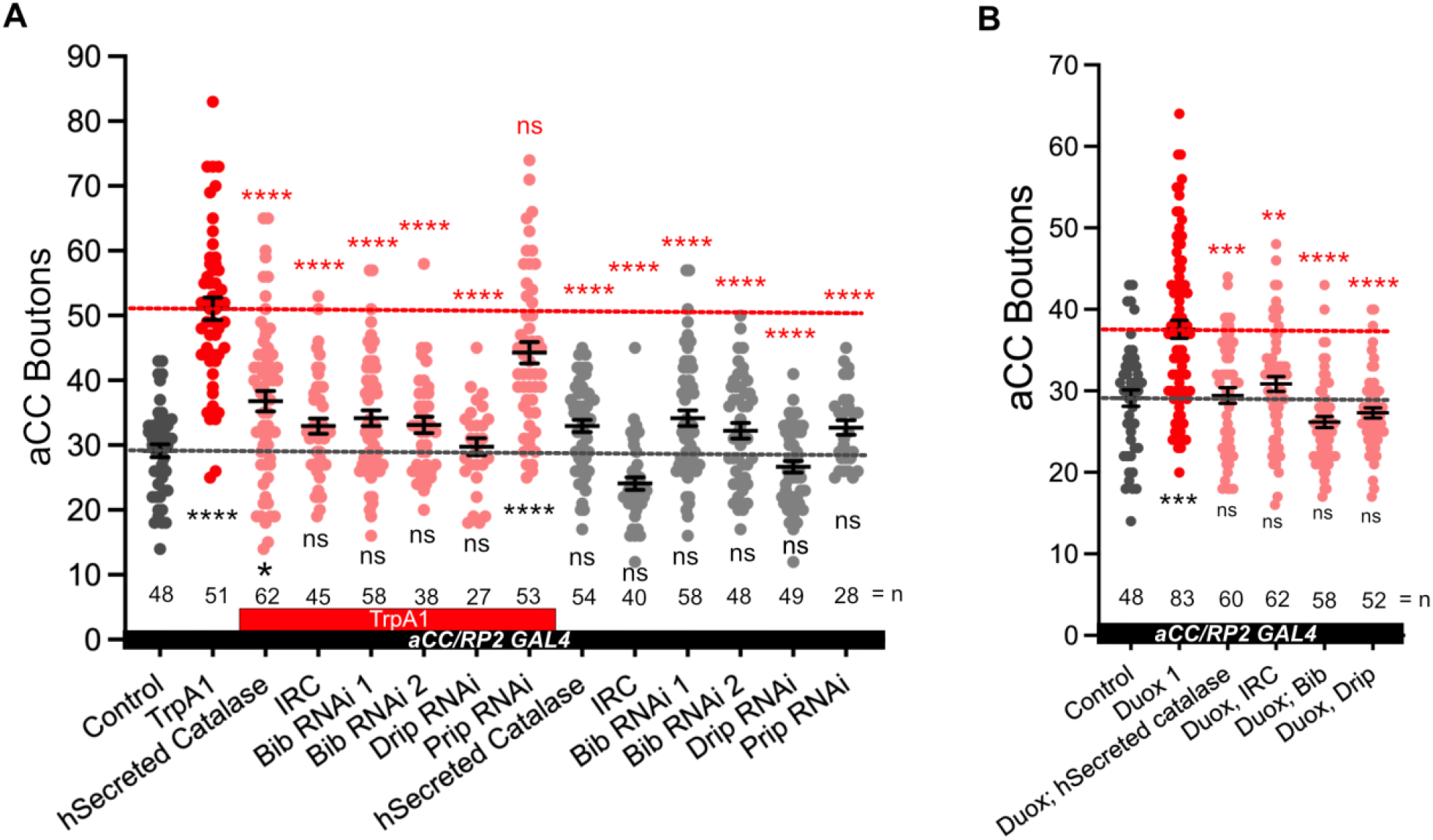
Aquaporins Bib and Drip are required for activity-regulated plasticity at the neuromuscular junction. A) Dot-plot quantification shows NMJ bouton number increases in response to cell-specific activity and the rescue of the phenotype when secreted catalases are expressed or aquaporins Bib or Drip are knock down. When the RNAi lines are represented by triangles, it indicates that it is in combination with dTrpA1, when the RNAi lines are represented by circles, it is without dTrpA1. B) Dot-plot quantification shows NMJ bouton number increases in response Duox and the rescue of the phenotype when secreted catalases are expressed or aquaporins Bib or Drip are knock down. Mean ± SEM, Kruskal-Wallis test, *p<0.05, ** p<0.01, *** p<0.001, **** p<0.0001. Comparisons are made with the TrpA1 group in A and with Duox group in B.

Because NAPDH oxidases generate ROS extracellularly, we wanted to explore how extra-cellular ROS might enter the cell so as to act on intracellular signalling pathways that would regulate NMJ growth. Several studies, including one from this lab, have postulated a role for aquaporin channels, specifically those encoded by the genes *bib* and *Drip* (Albertini and Bianche, 2010; Dhawan et al., 2020; Dutta and Das, 2022). Indeed, for the presynaptic NMJ, we found that co-expression of *UAS-RNAi* constructs designed to knock down *bib* or *Drip*, but not those for *prip*, rescue NMJ growth phenotypes caused by dTRPA1-mediated overactivation. Expression of the *UAS-RNAi* constructs alone had no significant effect (Figure 3A). To further test the model that extracellular ROS generated by NADH oxidases cause structural change at the NMJ, we overexpressed Duox in aCC motoneurons and at the same time co-expressed *UAS-RNAi* constructs designed to knock down the aquaporin channel proteins Bib or Drip. In those neurons the Duox gain-of-function NMJ growth phenotype is fully rescued (Figure 3B).

In summary, our observations suggest that at the presynaptic NMJ, neuronal overactivation leads to activation of both NADPH oxidases, Duox and Nox, at the plasma membrane. These enzymes generate ROS at the extracellular face, which are then brought into the cytoplasm by aquaporin channels comprising Bib and Drip. Inside the cell, the ROS act on intracellular membrane-localised signalling pathways that regulate synaptic terminal structure and size, including the phosphatase PTEN and DJ-1ß, as previously shown (Figure 4) (Oswald et al., 2018).

**Figure 4.**
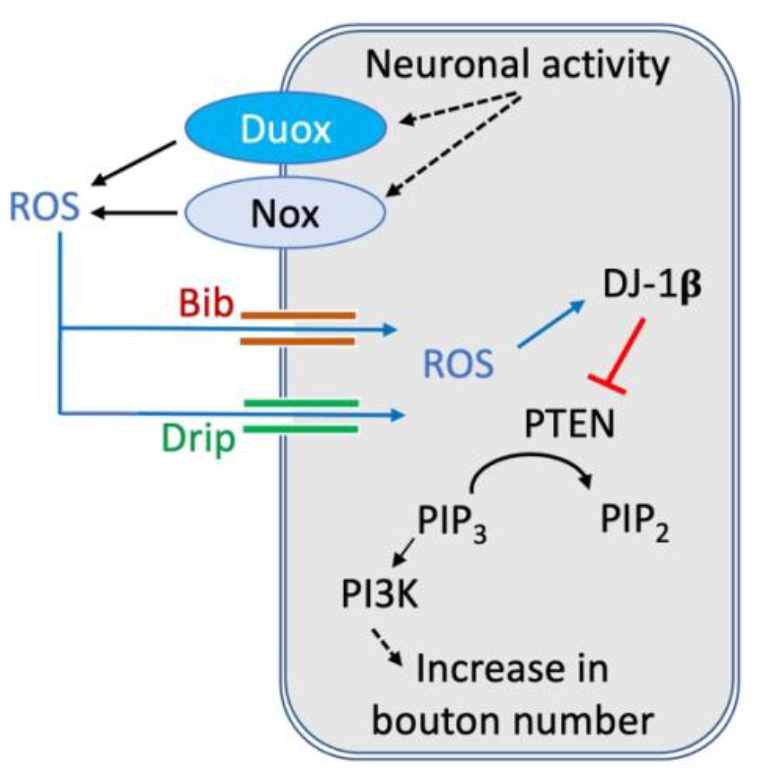
Model of activity-regulated plasticity at the neuromuscular junction mediated by ROS signalling. Neuronal activity leads to activation of both calcium regulated NADPH oxidases, Duox and Nox, at the presynaptic motoneuron terminal. Extracellularly generated ROS reintroduced into the presynaptic cytoplasm via aquaporin channels, including Bib and Drip. ROS modulated PI3Kinase signalling at the plasma membrane via oxidation of DJ-1ß, which when oxidised increases binding and inhibition of the PTEN phosphatase, thus causing increased PI3Kinase signalling activity, stimulating growth and addition of synaptic release sites.

## 3 Discussion

ROS have increasingly been recognised as signalling molecules required for nervous system development and function, from regulating the dynamics of the growth cone cytoskeleton to synaptic transmission and learning (see Terzi and Suter, 2020). At the Drosophila NMJ, ROS have been shown necessary for activity-induced synaptic terminal growth (Oswald et al., 2018). ROS have also been shown causative and sufficient to induce changes at synaptic terminals; when accumulating as a result of physiological dysfunction, leading to oxidative stress (Milton et al., 2011), or following manipulations that increase ROS levels (Milton et al., 2011; Hussain et al., 2018; Peng et al. 2019). While mitochondria are a major source of cellular ROS (Murphy, 2009; Zorov et al., 2014; Sanz, 2016), it has remained unclear to what extent mitochondrial ROS directly impact on events at the plasma membrane, such as modulation of PTEN-PI3Kinase signalling, which regulates synaptic terminal growth (Acebes, et al., 2012; Jordán-Álvarez et al., 2012; Martín-Peña et al., 2006; Oswald et al, 2018), or oxidation of ion channel subunits that modulate neuronal excitability (Kempf et al., 2019).

## Differential requirements for NADPH oxidases in pre- vs postsynaptic compartments

In this study we focused on NADPH oxidases as generators of ROS that are ideally positioned to influence signalling at the plasma membrane. Working with the NMJ in the Drosophila larva as an experimental *in vivo* model system, we demonstrated that both NADPH oxidases, Nox and Duox, are required for activity-induced growth (Figure 1). Both enzymes are endowed with N-terminal calcium binding EF-hand motifs, linking their activity to intracellular calcium levels, as shown for Drosophila Duox (Ha et al; 2009; Rigutto et al., 2009; Razzell et al., 2013) and the vertebrate homologue, Nox5 (Bánfi et al., 2004; Millana et al., 2020). Conversely, over-expression of either enzyme is sufficient to phenocopy such presynaptic terminal growth (Figure 2). Curiously, the requirement for NADPH oxidases in regulating dendritic growth is different, with only Duox, but not Nox, mediating activity-induced reduction of dendritic arbor size (Dhawan et al., 2020). This difference in pre- *versus* postsynaptic compartment regulation is mirrored by their differential sub-cellular localisation, with tagged Nox protein being effectively excluded from the postsynaptic dendritic arbors, unlike Duox (Figure 2C). Apart from this differential requirement in pre- *versus* postsysnaptic compartments, it is unclear to what extent Nox and Duox might perform different functions during activity-induced growth. At the NMJ, where both are present and required, we found no difference in phenotypes following RNAi knockdown or mis-expression. Curiously, phenotypes were also comparable regardless of whether the expression of both enzymes was manipulated simultaneously or individually, suggesting either a saturation of phenotype or, speculatively, that Nox and Duox might operate in the same signalling pathway with their activation contingent on one another.

### NADPH oxidases generate extracellular ROS and mediate autocrine signalling

Because Nox and Duox generate ROS at the extracellular face they have the potential for inter-cellular signalling, for example as documented during wound healing (Razzell et al, 2013; Niethammer et al., 2009, 2016). Indeed, within the dense meshwork of neuronal processes and synapses of the CNS, we recently found that reduction of extracellular hydrogen peroxide in the immediate vicinity of dendritic processes (by mis-expression of a secreted catalase) or attenuation of ROS entry into those dendrites (by knock-down of aquaporins), both cause significant dendritic over-growth (Dhawan et al., 2020). This suggests that within the densely innervated central neuropile, extracellular ROS generated, including those from activity-regulated NADPH oxidases, might function as local signals to which neurons respond with adjustments of their synaptic terminals. This contrasts with the peripheral Drosophila larval NMJ, where we did not see any significant changes in synaptic terminal morphology following manipulations that would either reduce entry of ROS into the presynaptic terminal or reductions of extracellular ROS (Figure 3). These observations suggest that at the presynaptic NMJ, NADPH oxidases might be required only under conditions of elevated neuronal activity. While these data further suggest that at the presynaptic NMJ, NADPH oxidase-generated ROS are principally engaged in autocrine signalling, we cannot currently exclude the potential for inter-cellular signalling to adjacent muscles and glia.

Autocrine ROS signalling at both pre- and postsynaptic compartments is underlined by the requirement for the aquaporin channel proteins, Bib and Drip (Figure 3) (Dhawan et al., 2020). Some studies have questioned the extent to which Bib might function as an aquaporin, as unable to form effective water channels in a heterologous expression system (Tatsumi et al., 2009; Kourghi et al., 2017). However, in this and in a pervious study (Dhawan et al., 2020), Bib RNAi knockdown produces synaptic terminal phenotypes indistinguishable from knockdown of Drip, or from mis-expression of secreted forms of catalase (Dhawan et al., 2020). This suggests that Bib functions in the same pathway as the aquaporin Drip, potentially forming part of a heteromeric channel with permeability for hydrogen peroxide.

### Independent, local regulation of pre- and postsynaptic terminal growth

Over-activation of neurons results in changes to both pre- and postsynaptic terminals, though it has been unclear in how far such changes in growth of input and output compartments might be co-ordinately regulated. Working with this experimental system we happen to have identified two sets of manipulations that suggest the growth of pre- and postsynaptic terminals can be regulated independently of each other. First, in motoneurons that have been over-activated by mis-expression of dTRPA1, RNAi knockdown of Nox has no effect on the activity-induced reduction of the postsynaptic dendrites, which receive all synaptic input from pre-motor interneurons (as Nox protein appears to be excluded from dendrites); yet at the output compartment, the presynaptic NMJ, of those same neurons, activity-linked overgrowth is significantly suppressed by knockdown of Nox. This contrasts with the effect of Duox knockdown under conditions of neuronal over-activation, with Duox RNAi suppressing over-activation phenotypes effectively at both pre- and postsynaptic terminals.

Second, RNAi knockdown alone of the genes coding for aquaporin channel proteins Bib or Drip cause significant dendritic overgrowth, without affecting the presynaptic NMJ. These manipulations suggest that, at least in Drosophila larval motoneurons, synaptic terminal growth can be regulated locally through ROS signalling, such that pre- and postsynaptic compartments can adjust independently from each other. This makes sense when viewing extracellular ROS as local signals for over-activation, to which cells respond by adjusting their synaptic terminals. In this context, it remains to be seen to what extent extracellular ROS might impact on the regulation of synaptic transmission.

In summary, it is increasingly appreciated that ROS are important signals, whose signalling capability is proportional to the spatiotemporal precision attained. Sub-cellular specificity of ROS generators, such as the NAPDH oxidases studied here, is an important facet.

## 4 Materials and Methods

### Fly genetics

*Drosophila melanogaster* strains were maintained on a standard apple juice-based agar medium at 25°C. The following fly strains were used: *OregonR* (#2376 Bloomington Drosophila Stock Center), *UAS-dTrpA1* in attP16 (Hamada et al., 2008; FBtp0089791), *UAS-Duox.RNAi* (1) (#32903 BDSC; FBtp0064955), *UAS-Duox.RNAi* (2) (#38916 BDSC; FBgn0283531), *UAS-Nox.RNAi* (1) (Ha et al., 2005b; FBal0191562), *UAS-Nox.RNAi* (2) (#32433 BDSC; FBgn0085428), *UAS-bib.RNAi* (1) (#57493 BDSC; FBtp0096443), *UAS-bib.RNAi* (2) (#27691 BDSC; FBtp0052515), *UAS-Drip.RNAi* (1) (#44661 BDSC; FBtp0090566), *UAS-Drip.RNAi* (2) (#106911 Vienna Drosophila Resource Centre; FBtp0045814) (Begland et al., 2012), *UAS-Prip.RNAi* (2) (#44464 BDSC; FBtp0090258), *UAS-Duox* (1) (Ha et al., 2005b), *UAS-Duox∷mRuby2∷HA* (2) (this paper), *UAS-Nox∷YPet* (this paper), *UAS-hCatS (human secreted catalase)* (FBal0190351; Ha et al., 2005b; Fogarty et al., 2016), *UAS-extracellular immune-regulated catalase* (Irc) (FBal0191070, Ha et al., 2005b).

Transgene expression was carried out at 25°C, unless otherwise noted, targeted to RP2 and aCC motoneurons using the following Gal4 expression line: *RN2-O-Gal4*, *UAS-FLP, tubulin84b-FRT-CD2-FRT-Gal4; RRFa-Gal4, 20xUAS-6XmCherry∷HA* (Pignoni and Zipursky, 1997; Fujioka et al., 2003; Shearin et al., 20014). Briefly, RN2-GAL4 expression in RP2 and aCC motoneurons is restricted to the embryo, but is maintained subsequently by FLPase-gated *tubulin84B-FRT-CD2-FRT-GAL4* (Ou et al., 2008). mCherry∷HA was used as morphological reporter. To study the localisation of the tagged *Nox∷YPet* and *Duox∷mRuby2∷HA* transgene expression was targeted to aCC motoneurons using the *GMR94G06-Gal4* (#40701 BDSC; FBgn0053512; Pérez-Moreno and O’Kane, 2019). *pJFRC12-10XUAS-IVS-Nox-YPet* (GenBank OP716753) in landing site VK00040 [cytogenetic location 87B10] was generated by Klenow assembly cloning (tinyurl.com/4r99uv8m). Briefly, from pJFRC12-10XUAS-IVS-myr-GFP plasmid DNA we removed the coding sequence for *myr∷GFP* using XhoI and XbaI, and replaced it with *Nox* cDNA from DGRC clone FI15205 (pOTB7 vector backbone; kindly provided by Kenneth H. Wan, DGRC Stock 1661239; https://dgrc.bio.indiana.edu//stock/1661239 ; RRID:DGRC_1661239), its 3’ stop codon replaced by a flexible glycine-serine linker, followed by YPet (Nguyen and Daugherty, 2005). Similarly, we created *pJFRC12-10XUAS-IVS-Duox-mRuby2-HA* (GenBank OP716753) in landing site VK00022 [cytogenetic position 57A5] using *Duox* cDNA kindly provided by Won-Jae Lee, its 3’ stop codon replaced by a flexible glycine-serine linker, followed by mRuby2 (Lam et al, 2012), followed by another glycine-serine flexible linker and four tandem repeats of the hemagglutinin (HA) epitope. Transgenics were generated via phiC31 integrase-mediated recombination (Bischof et al.; 2007) into defined landing sites by the FlyORF Injection Service (Zürich, Switzerland).

### Dissections and immunocytochemistry

Flies were allowed to lay eggs on apple-juice agar based medium overnight at 25 °C. Larvae were then reared at 25°C on yeast paste, while avoiding over-crowding. Precise staging of the late wandering third instar stage was achieved by: a) checking that a proportion of animals from the same time-restricted egg lay had initiated pupariation; b) larvae had reached a certain size and c) showed gut-clearance of food (yeast paste supplemented with Bromophenol Blue Sodium Salt (Sigma-Aldrich)). Larvae were dissected in Sorensen’s saline, fixed for 5 min at room temperature in Bouins fixative or 10 min paraformaldehyde (Agar Scientific) when staining for GFP/YPet epitopes, as previously detailed (Oswald et al., 2018). Wash solution was Sorensen’s saline containing 0.3% Triton X-100 (Sigma-Aldrich) and 0.25% BSA (Sigma-Aldrich). Primary antibodies, incubated overnight at 10°C, were: Goat-anti-HRP Alexa Fluor 488 (1:1000) (Jackson ImmunoResearch Cat. No. 123-545-021), Rabbit-anti-dsRed (1:1000) (ClonTech Cat. No. 632496), Mouse nc82 (Bruchpilot; Developmental Studies Hybridoma Bank Cat No nc82), Chicken anti-GFP (1:5000) (abcam Cat No ab13970); secondary antibodies, 2 hr at room temperature: Donkey anti-mouse Alexa Fluor 647; Donkey-anti-Rabbit CF568 (1:1200) (Biotium Cat. No. 20098), Donkey anti-Chicken CF488 (1:1000) (Cambridge Bioscience Cat No 20166) and goat anti-Rabbit Atto594 (1:1000) (Sigma-Aldrich Cat No 77671-1ML-F). Specimens were cleared in 70% glycerol, overnight at 4°C, then mounted in Mowiol.

### Image acquisition and analysis

Specimens were imaged using a Leica SP5 point-scanning confocal, and a 63x/1.3 N.A. (Leica) glycerol immersion objective lens and LAS AF (Leica Application Suite Advanced Fluorescence) software. Confocal images were processed using ImageJ (to quantify active zones) and Affinity Photo (Adobe; to prepare figures). Bouton number of the NMJ on muscle DA1 from segments A3-A5 was determined by counting every distinct spherical varicosity along the NMJ branch.

To study if genetic manipulations targeted to aCC and RP2 motoneurons change muscle size we measured surface area of DA1 muscles, imaged under DIC optics using a Zeiss Axiophot compound microscope and a Zeiss Plan-Neofluar 10x/0.3 N.A. objective lens. Images were taken with an Orca CCD camera (Hamamatsu) and muscle surface area was determined using ImageJ by multiplying muscle length by width. Quantification of key representative experiments, covering most transgenic lines used and conditions where genetic manipulation of aCC motoneurons cause significant changes in bouton number, show no statistically significant differences in average muscle size, which is used as an indicator of overall animal size. Correlation between individual muscle sizes and bouton numbers show that the biggest differences in muscle surface area is due to dissection artefact of differences to the extent that larval filets are stretched, rather than differences in animal or muscle growth, which would lead to clear correlations between measured muscle surface area and NMJ bouton number (see supplementary figure 2). Taking account of this, bouton numbers are shown as raw counts, not normalized to average muscle surface area.

Representative schematics, drawings and plates of photomicrographs were generated with Affinity Photo (Serif Ltd., United Kingdom).

### Statistical analysis

All data handling was performed using Prism software (GraphPad). NMJ bouton number data was tested for normal/Gaussian distribution using the D’Agostino-Pearson omnibus normality test. When normal distribution was confirmed the statistically comparisons were done using one-way analysis of variance (ANOVA), with Tukey’s multiple comparisons test. When non-normal distribution was confirmed the statistically comparisons were done using Kruskal-Wallis test.

## 6 Supplementary Figures

**Supplementary Figure 1.**
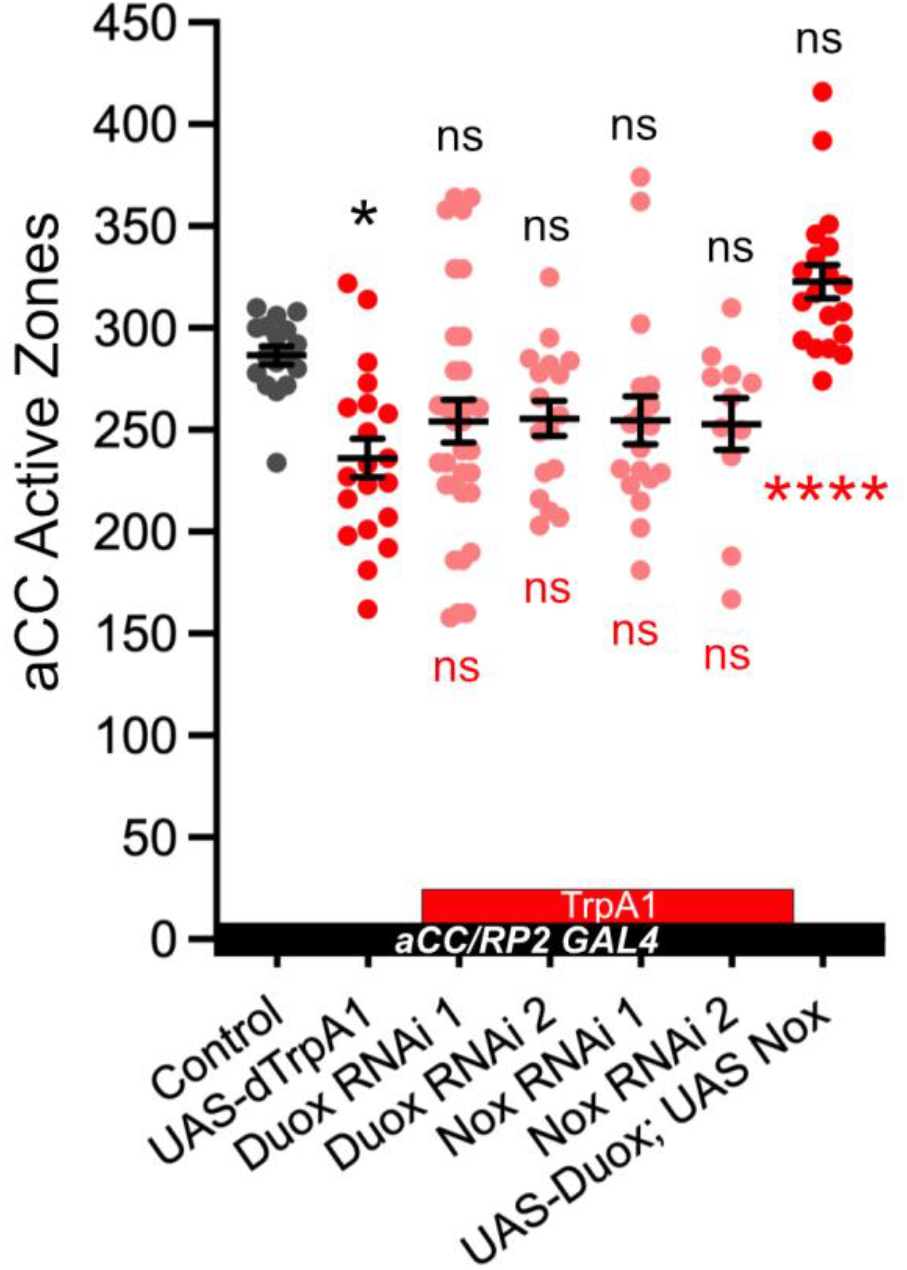
NADPH oxidases, dDuox and dNox, are not necessary for activity-regulated synaptic number at the neuromuscular junction. Dot-plot quantification shows active zone number increases following overactivation (dTrpA), but there are not significant differences when manipulating NADPH oxidases. Mean ± SEM, ANOVA, * p<0.05, **** p<0.0001.

**Supplementary Figure 2.**
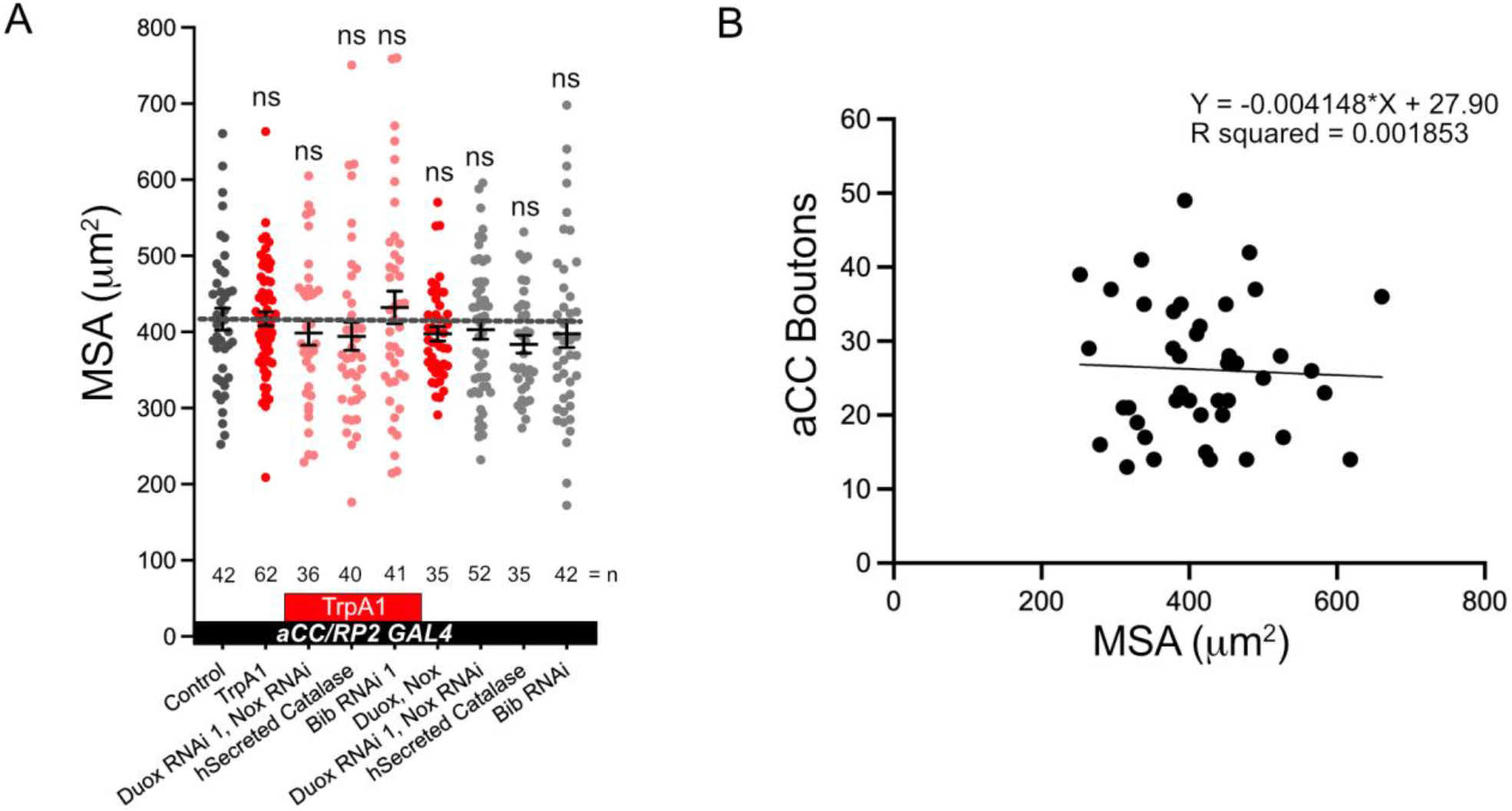
Muscles size. A) Dot-plot quantification shows no statistically significant differences between genotypes in average muscle surface area (MSA). Mean ± SEM, Kruskal-Wallis test. B) Linear regression using the control data shows not correlation between aCC NMJ terminal bouton numbers and muscle size, p-value = 0.7866.

## 7 Author Contributions

D.S.C, M.C.W.O, A.M. and M.L. conceived of the study and wrote the manuscript. D.M.D.B. cloned Duox and Nox transgenes. M.L. generated transgenic stocks. D.S.C. and M.C.W.O. carried out all experiments and analysed data.

## 8 Funding

This work was made possible through support by the Biotechnology and Biological Sciences Research Council (BBSRC) to M.L. (BB/R016666/1 and BB/V014943/1). D.S.-C. was supported by the European Molecular Biology Organization (EMBO) with a long-term EMBO fellowship (ALTF 62-2021) and a John Stanley Gardiner studentship to A.M. The work benefited from the Imaging Facility, Department of Zoology, supported by Matt Wayland and funds from a Wellcome Trust Equipment Grant (WT079204) with contributions by the Sir Isaac Newton Trust in Cambridge, including Research Grant [18.07ii(c)].

## 9 Acknowledgments

The authors would like to thank Niklas Krick for feedback on the manuscript. The authors are grateful to Andreas Bergmann, Paul Garrity, Won-Jae Lee, Paul Martin, Sean Sweeney, Helen Weavers, and Will Wood, as well as the Bloomington Drosophila Stock Center and Vienna Drosophila Resource Centre for generously providing fly stocks; and to Won-Jae Lee for providing DNA containing Duox cDNA, and the Drosophila Genomics Resource Center (DGRC), supported by NIH grant 2P40OD010949, for clone FI15205 containing Nox cDNA.

## Notes

### Competing Interest Statement

The authors have declared no competing interest.

